# Male-biased recombination at chromosome ends in a songbird revealed by precisely mapping crossover positions

**DOI:** 10.1101/2023.12.19.572321

**Authors:** Hongkai Zhang, Max Lundberg, Suvi Ponnikas, Dennis Hasselquist, Bengt Hansson

## Abstract

Recombination plays a crucial role in evolution by generating novel haplotypes and disrupting linkage between genes, thereby enhancing the efficiency of selection. Here, we analyse the genomes of twelve great reed warblers (*Acrocephalus arundinaceus*) in a three-generation pedigree to identify precise crossover positions along the chromosomes. We located more than 200 crossovers and found that these were highly concentrated towards the telomeric ends of the chromosomes. While the number of recombination events was similar between the sexes, the crossovers were located significantly closer to the ends of paternal compared to maternal chromosomes. The frequency of crossovers was similar between intergenic and genic regions, but within genes, they occurred more frequently in exons than in introns. In conclusion, our study of the great reed warbler revealed substantial variation in crossover frequencies within chromosomes, with a distinct bias towards the sub-telomeric regions, particularly on the paternal side. These findings emphasise the importance of thoroughly screening the entire length of chromosomes to characterise the recombination landscape and uncover potential sex-biases in recombination.

**Article summary:** The genetic exchange between the paternal and maternal chromosomes during meiosis – recombination – plays a crucial role in evolution by generating new haplotypes that natural selection can act upon. By analysing genomic data of a three-generation family of great reed warblers, we detected precise locations of approximately 200 recombination events in the genome of these birds. This unveiled a prominent sex-bias with recombination occurring more often towards chromosome ends in males than in females.

## 1 Introduction

Recombination has profound evolutionary implications by generating new haplotypes that natural selection can act upon. The process of reshuffling haplotypes through recombination breaks linkage disequilibrium and reduces the interference between linked loci, which otherwise limits the action of natural selection (Felsenstein 1974; Otto 2021). Recombination disconnects beneficial and deleterious alleles at linked loci, facilitating adaptive evolution (increasing the frequency of advantageous alleles) and purging the genetic load (reducing the frequency of deleterious alleles). The Y and W chromosomes of mammals and birds illustrate the importance of recombination, as their prolonged periods without recombination have resulted in significant degeneration and paucity of genes (Charlesworth and Charlesworth 2000; Bachtrog 2013).

Recombination not only has evolutionary implications but can also be subject to selection and undergo evolutionary changes itself. Studies have demonstrated variation in recombination rates across clades, species, populations, and between sexes (Ritz et al. 2017; Stapley et al. 2017; Peñalba and Wolf 2020). For example, fungi generally exhibit higher rates of recombination (averaging 48.7 cM/Mb) compared to plants (averaging 1.9 cM/Mb) (Stapley et al. 2017), and among mammals, recombination rates range from 0.2 cM/Mb in opossums to 1.6 cM/Mb in dogs (Dumont and Payseur 2008). The variation in recombination rate across species and deeper lineages can be attributed, at least in part, to evolved differences in chromosome size and number. The presence of more and smaller chromosomes tends to increase recombination rates since each chromosome (or chromosome arm) requires a minimum of one crossover (CO) on one of the two sister chromatids (Coop and Przeworski 2007; Stapley et al. 2017). Regarding sex differences, the most extreme scenario is when recombination is entirely absent in one sex (achiasmy). This lack of recombination coincides almost exclusively with the heterogametic sex, such as male *Drosophila* (XY) and female Lepidoptera (ZW) (Haldane 1922; Huxley 1928; John et al. 2016). However, in many plants and animals, the degree of sexual dimorphism in recombination (heterochiasmy) is less pronounced, with at least some level of recombination in both sexes (Burt et al. 1991; Broman et al. 1998; Maddox et al. 2001; Berset-Brändli et al. 2008; Wellenreuther et al. 2013; Bergero et al. 2019; Malinovskaya et al. 2020). Generally, it is not surprising that recombination can evolve since it exhibits typical features of evolving traits, such as variation among individuals and a heritable component, as seen in humans (Kong et al. 2004) and sheep (Johnston et al. 2016). In fact, evidence from experimental populations in *Drosophila* shows that the rate of recombination can be manipulated over short evolutionary time scales (Aggarwal et al. 2015; Kohl and Singh 2018).

Quantifying the recombination landscape – the local recombination rate variation along the chromosomes – can provide valuable insights into the influence of recombination on various biological processes. For instance, as selection operates on chromosome regions (linked selection), low recombining regions often exhibit reduced selection efficiency at single mutations, lower local effective population size (N_e_), and stronger genetic drift (Peñalba and Wolf 2020). Moreover, recombination can contribute to elevated GC content through GC-biased gene conversion (Mugal et al. 2015) and increase genetic variation by locally raising the mutation rate (Filatov and Gerrard 2003; Hellmann et al. 2003; Huang et al. 2005; Arbeithuber et al. 2015; Halldorsson et al. 2019). As expected, the local recombination rate correlates with various population genetic parameters, including linkage disequilibrium, nucleotide diversity, GC and repeat content, and gene density (Peñalba and Wolf 2020; Ponnikas et al. 2022). A common pattern observed in many species is an increase in recombination rates towards the telomeres and decreasing rates around the centromeres (Vincenten et al. 2015; Limborg et al. 2016; Haenel et al. 2018; Sardell and Kirkpatrick 2020). Mechanistically, heterochromatin, which is enriched around the centromeres, can prevent local recombination by hindering polymerase accessibility or through repression of double-strand break (DSB) formation caused by methylation, RNAi or specific enzymes (Ellermeier et al. 2010; Melamed-Bessudo and Levy 2012; Wijnker et al. 2013). Quantifying the recombination landscape allows the identication of recombination hotspots where recombination is particularly frequent, as well as cold spots, where recombination is infrequent. Notably, detailed analysis of recombination hotspots in humans and mice led to the discovery of the *PRDM9* gene as a key regulator of the local recombination rate (Myers et al. 2008; Baudat et al. 2010). However, while *PRDM9* is present in most mammals, it is absent in birds, suggesting different mechanisms of recombination regulation across taxa (Singhal et al. 2015). Furthermore, there are still many unanswered questions, such as why and how the recombination landscapes sometimes differ between sexes, given the observation of higher recombination rates closer to the chromosome ends in males and more uniform recombination landscapes in females (Sardell and Kirkpatrick 2020). Exploring these differences and understanding their underlying mechanisms remain areas of active research.

The initial data of recombination came from direct observations of chiasmata and from segregation of Mendelian traits in model organisms, such as *Drosophila* (Morgan 1910; Haldane 1920; Morton et al. 1976). Subsequently, segregation analysis of molecular markers in large pedigrees became popular for constructing linkage maps and inferring recombination distances in centimorgans (cM) (Robinson 1996; Dumont et al. 2011). Combining physical maps and linkage maps allowed for the inference of local recombination rates along chromosomes (Groenen et al. 2009; Backström et al. 2010; Kawakami et al. 2014; Johnston et al. 2017). The advent of next-generation sequencing (NGS) enabled the screening of dense marker sets in many individuals, facilitating pedigree-free methods to study recombination rates. These methods involve assessing the local level of linkage disequilibrium (LD) and assuming a negative correlation between LD and recombination rate (Singhal et al. 2015; Provost et al. 2022). This has been accompanied by progress in statistics and model inferences of the population-level recombination landscape (McVean and Auton 2007; Chan et al. 2012; Gao et al. 2016; Adrion et al. 2020). Furthermore, dense marker data provided by NGS allows locating individual crossover positions with high precision on chromosomes by phasing alleles from their segregation pattern in small pedigrees (Smeds et al. 2016), or, as applied most recently, by directly generating haploid data through sperm sequencing (Bell et al. 2020). In birds, high-resolution recombination positioning using NGS-generated single nucleotide polymorphism (SNP) data has so far been applied to a collared flycatcher (*Ficedula albicollis*) three-generation pedigree (Smeds et al. 2016).

In this study, we aim to identify recombination positions at a highly detailed chromosomal scale by analysing genome-wide SNPs segregating in a three-generation pedigree of the great reed warbler (*Acrocephalus arundinaceus*). To achieve this, we sequenced the genomes of the individuals in the pedigree, mapped the reads to the reference genome of the great reed warbler (Sigeman et al. 2021), and called SNPs. Then, we employed the newly developed *RecView* R package (Zhang and Hansson 2023) to analyse the genome-wide SNP data, allowing us to examine the segregation patterns at each SNP in the pedigree and pinpoint recombination positions between genomic regions inherited from different grandparents. Previous studies have indicated significant heterochiasmy in the great reed warbler, with females exhibiting nearly twice the recombination rate compared to males (Hansson et al. 2005; Dawson et al. 2007). However, these earlier studies were limited by the use of few markers covering only small portions of the chromosomes. In contrast, our present study utilises millions of SNPs, which enables us to examine whether recombination events preferentially occur in specific regions of the chromosomes and whether the number and positions of recombination events differ between paternal and maternal chromosomes.

## 2 Materials and methods

### 2.1 Generating the SNP dataset

The great reed warbler is a large Acrocephalid warbler and long-distance migrant that spends the winter in sub-Saharan Africa, and returns to breed in reed lakes in Europe and western Asia during the summer (Helbig and Seibold 1999; Lemke et al. 2013; Koleček et al. 2016; Sjöberg et al. 2021). We selected a three-generation pedigree from our long-term study population of great reed warblers at Lake Kvismaren, Sweden (Bensch et al. 1998; Hasselquist 1998; Tarka et al. 2014; Hansson et al. 2018). The pedigree consisted of 12 individuals (**Figure 1**) and included four grandparents (F0 generation), two parents (F1 generation) and six offspring (F2 generation), with the offspring belonging to the 1998 cohort (five males and one female; **Table S1**).

**Figure 1.**
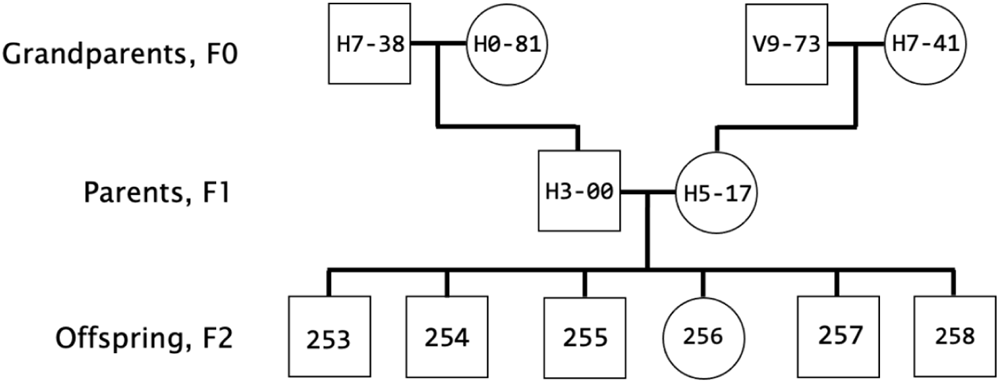
The three-generation great reed warbler pedigree analysed in the present study. Project-specific individual codes are given. Squares show males, circles females.

We used a phenol-chloroform protocol to extract genomic DNA from blood (stored in SET buffer) from each of the 12 individuals. Sequencing libraries were created using the TruSeq (Illumina) protocol with 350 bp insert size, and sequenced on a NovaSeq 6000 (Illumina) using a 2×150 bp setup and targeting 50x coverage. Libraries and sequencing were performed by SciLifeLab, Uppsala, Sweden.

The raw sequence reads were trimmed with *trimmomatic* version 0.39 (Bolger et al. 2014), mapped to the reference genome assembly of the great reed warbler (Sigeman et al. 2021) using *bwa mem* version 0.7.17 (Li 2013), and read duplicates were removed with *PicardTools* version 2.27.5 (Broad Institute). Then, variants were called with *freebayes* version 1.3.2 (Garrison and Marth 2012), producing a VCF file. Variants in annotated repeat intervals were removed using *vcftools* version 0.1.16 (Danecek et al. 2011). Only bi-allelic variants were kept, and decomposed for complex and multinucleotide variants, using *vcftools* and *VT decompose_blocksub* version 0.5 (Tan et al. 2015). After indels had been removed using *vcffilter* from *vcflib* version 2017-04-04 (Garrison et al. 2022), the SNPs were divided into an autosome and a Z-linked dataset. Filtering on quality, strandedness, read placement and genotype coverage (with -f “QUAL > 30 & SAF > 0 & SAR > 0 & RPR > 0 & RPL > 0” and -g “DP > 9” in *vcffilter*) were applied for both datasets. Finally, SNPs with missing data were removed using *vcftools*.

### 2.2 Chromosome-level assembly and chromosome arms

The genome assembly of the great reed warbler consists of relatively few large scaffolds (Sigeman et al. 2021). Some of the smaller chromosomes are represented by a single scaffold (chromosomes 4A, 13, 14, 19, 20, 23 and 30; Sigeman et al. 2021), and the Z chromosome (which includes a translocated part of chromosome 4A) has previously been assembled to chromosome-level (Ponnikas et al. 2022). The remaining chromosomes are represented by between two and nine scaffolds (Sigeman et al. 2021). To ordered and oriented the scaffolds of the remaining chromosomes, we used synteny analyses to several sources. We used the chromosome-level assemblies of the zebra finch (*Taeniopygia guttata*; bTaeGut1.4.pri, NCBI BioProject ID PRJNA489098) and the great tit (*Parus major*; Parus_major1.1, NCBI BioProject ID PRJNA208335) to assign scaffolds to chromosomes. Next, the scaffolds were ordered and oriented using *de novo* assemblies of two additional great reed warbler individuals (one male and one female; neither being the individual used for the original assembly; constructed with 10X linked read and HiC data) (B. Hansson *et al*., unpubl. data) and the chromosome-level assembly of another *Acrocephalus* species, the Eurasian reed warbler (*A. scirpaceus*; Sætre et al. 2021). During this process, we detected that the scaffold Contig1 on chromosome 2 was wrongly assembled, and this was corrected (**Table S2**). When the two *de novo* assemblies gave contradictory suggestions about the order or orientation (which could happen for methodological and biological reasons, such as the presence of inversion polymorphisms), we selected the order or orientation that was supported by the Eurasian reed warbler assembly and/or the recombination analysis in this study (**Table S2**). We did not manage to include some of the micro-chromosomes and chromosome 16 (the latter contains the structurally complicated major histcompatibility complex; Westerdahl et al. 2022). Moreover, one chromosome was deemed too short for reliable analysis of recombination (chromosome 30; 1.3 Mb) and one had several scaffolds with uncertain order and orientation (chromosome 22; see below). Thus, the final number of chromosomes assembled to chromosome-level and included in the present analysis was 29 autosomes and the Z chromosome (**Table S2**), which is similar to that of the great tit and Eurasian reed warbler genome assemblies, but lower compared to the zebra finch where all autosomes are (at least partly) assembled (bTaeGut1.4.pri; NCBI BioProject ID PRJNA489098). We named the chromosomes according to their zebra finch homologs (this study and Sigeman et al. 2021). The total length of the autosomal assembly was 978 Mb, which is similar to the great tit and zebra finch assemblies (1020 and 1056 Mb, respectively).

Autosomes 1, 1A, 2, 3 and 4 are sub-telocentric or sub-metacentric (*i.e.*, have two chromosome arms) in the zebra finch, whereas the remaining autosomes are telocentric (Knief and Forstmeier 2016). For these sub-telocentric or sub-metacentric chromosomes, we estimated the approximate location of the centromere based on low nucleotide diversity among the 12 sequenced individuals (extremely low nucleotide diversity is expected in the centromeric regions; Stump et al. 2005), and its location in the zebra finch (Knief and Forstmeier 2016), and devided them into two arms with the longer arm denoted q and the shorter p (**Table S3**). The Z chromosome is acrocentric in the zebra finch (Knief and Forstmeier 2016), but we did not divide this chromosome in the great reed warbler as there is an exceptionally low nucleotide diversity in a large central part of the great reed warbler Z chromosome (the region between c. 15-70 Mb; total length 87.5 Mb; Ponnikas et al. 2022). However, the Z chromosome has a small pseudoautosomal region (PAR), a region that recombines with the W chromosome in females. As we have one female offspring in our dataset, which has inherited a W chromosome from its mother, we present data for the PAR (1.1 Mb; where both sexes may recombine) and the non-PAR (86.4 Mb; where only males recombine) of the Z chromosome, separately.

### 2.3 Localising crossovers using the *RecView* R package

For viewing and locating crossovers (COs), we used *RecView*, an R package that we recently developed specifically for segregation analysis of SNPs in three-generation pedigrees (Zhang and Hansson 2023). *RecView* requires two input files, one providing the order and orientation of the scaffolds of the reference genome, and one providing the genotype data of the individuals. The genotype files for the autosomes and the Z chromosome were generated by extracting genotypes from the VCF files using *vcftools* (command: vcftools --gzvcf [vcf file] --extract-FORMAT-info GT --stdout > [output file]), and converting the data using the *make_012gt()* function in *RecView* (for a description of the input files, see Zhang and Hansson 2023).

In the *RecView* ShinyApp, we seleted the Cumulative Continuity Score (CCS) algorithm with a threshold of 50 (represented with CCS50) to locate the positions of putative COs (for a description of CCS, see Zhang and Hansson 2023). We conducted a manual examination of all CO positions and detected a few artefacts where identical CO positions occurred in more than one offspring and always coincided with scaffold borders. We strongly suspect that this indicates scaffolds that had been wrongly ordered or oriented, and therefore corrected and recalculated the CO positions accordingly (most of these were small and some of them were also supported by the Eurasian reed warbler assembly; corrected order and orientation are given in **Table S2**). However, chromosome 22 had two such problematic scaffolds (contig83 and contig39_split_69700), and since we could not order and orientation these scaffolds reliably, we excluded this chromosome from the analysis (**Table S2**). Thus, our analyses of CO positions included 34 autosomal arms (29 autosomes, five with two arms) and the PAR and non-PAR regions of the Z chromosome (**Table S4**).

### 2.4 Statistical tests

For each autosomal arm, we calculated both the sex-average and sex-specific recombination distance (cM) by dividing the number of paternal and maternal COs in the six offspring with the number of analysed meiosis (12 in total; one in each parent for each offspring) and multiplying the value with 100. The recombination distance in cM indicates the probability of observing a recombination event over a specific chromosome region (here, a chromosome arm). Next, the sex-average and sex-specific recombination distances were divided by the size of each autosomal arm to obtain respective recombination rates (cM/Mb). For the Z chromosome, the calculations of recombination distance and rate were the same as for the autosomes, but here we separated between the PAR where both parents may recombine and the rest of the chromosome where only the father may recombine.

We tested the difference between sexes in autosomal recombination distance (cM), and in autosomal recombination rate (cM/Mb), with paired t-tests on the autosomal arm-level using the *rstatix* R package (Kassambara 2022). Next, we tested whether CO positions differed between sexes by using both the actual and the proportional positions of COs on the chromosome arms (measured from the telomeric end of the chromosome arms) with a Mann-Whitney U test in the *stats* R package (R Core Team 2022).

We used available gene annotations (Sigeman et al. 2021) in *bedtools* (Quinlan and Hall 2010) to explore the overlap between autosomal COs and intergenic, genic, exon, intron, UTR and CDS regions. We focussed on the 6 Mb telomeric ends of the chromosomes because most COs (87%; see Results) were located within these chromosomal regions, and because this 6-Mb region differs from other parts of the chromosomes in, for example, gene density. We tested whether the number of autosomal COs differed between the following pairs of annotation features: intergenic vs genic regions, exon vs intron, and UTR vs CDS. Here we used chi-squared goodness-of-fit tests with the expected CO-numbers calculated from the total length (in bp) of the intervals of each feature within the 6 Mb regions, using the *rstatix* R package (Kassambara 2022).

We used *tidyverse* R package (Wickham et al. 2019) for data handling, and plotted the results with *ggplot2* (Wickham 2011).

## 3 Results

We evaluated 408 autosomal arms segregating in the pedigree (34 paternal and 34 maternal chromosome arms for each of the 6 offspring) and located 224 COs, of which 113 were paternal and 111 maternal (**Table S4**). For the Z chromosome, we identified 1 CO in the PAR (of maternal origin) and 6 COs on the remaining part of the Z where only males recombine (Table S4). The estimated precision of the recombination positions was generally high (mean = 557 bp; range 153-8333 bp). Of the 224 autosomal COs, 203 were single CO events on autosomal arms, 18 were double COs (*i.e.*, 9 arms had two COs), and three formed a triple CO event (*i.e.*, one arm had three COs; **Figure 2A**). This means that 213 autosomal arms (203+9+1) had at least 1 CO whereas 195 had no CO (**Figure 2A**). The number of autosomal arms with or without COs did not differ significantly (chi-square test; χ^2^ = 0.79, df = 1, p = 0.37). Most cases with more than one CO occurred on larger chromosome arms, and the shortest chromosome arm where this occurred was chromosome 15q (14.9 Mb) for which two COs were detected in two offspring (both originating from the maternal side; **Table S4**). The unique case with three COs on a chromosome arm (in a single offspring) occurred on the long arm (q) of chromosome 4, which is one of the macrochromosomes in songbirds (**Table S4**).

**Figure 2.**
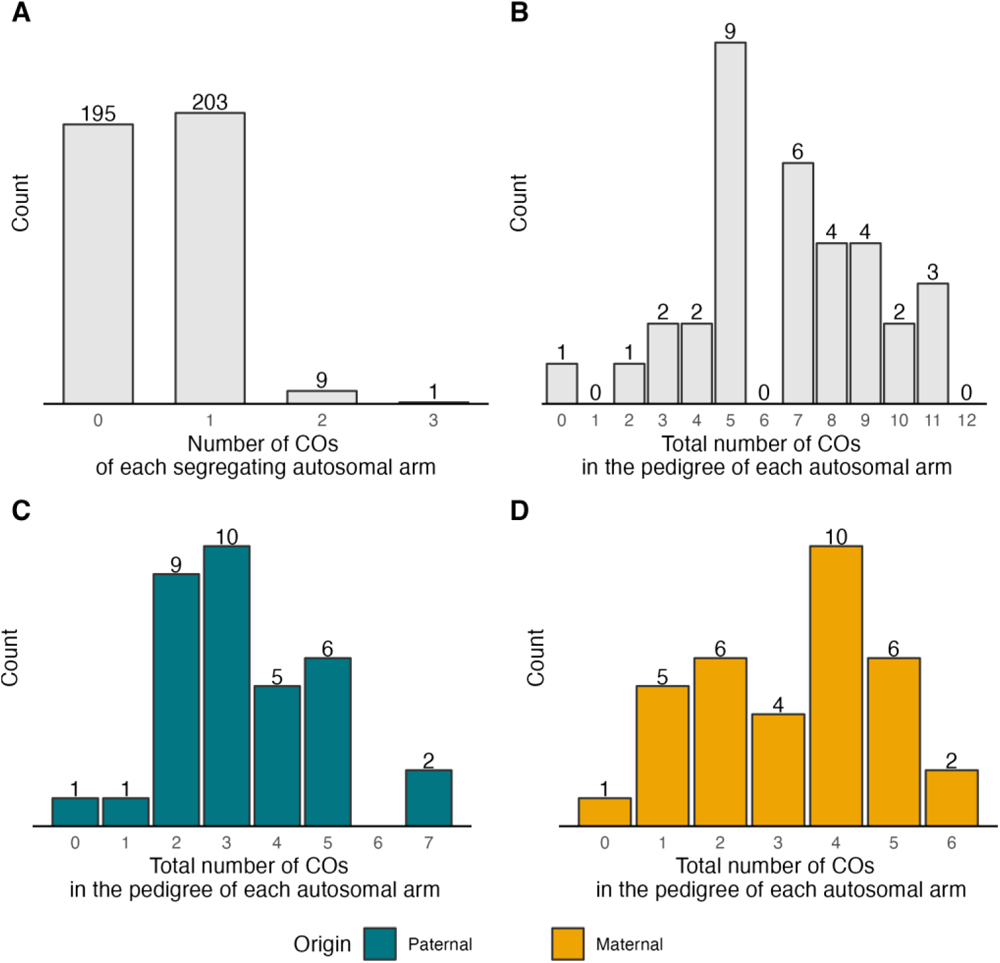
(A) The distribution of crossovers (COs) of each segregating autosomal arm. The total number of segregating autosomal arms evaluated was 408 (34 paternal and 34 maternal autosomal arms for each of the six offspring). (B-D) The distribution of the total number of COs in the pedigree of each autosomal arm (n = 34) considering COs of (B) both paternal and maternal origin, (C) only paternal origin, and (D) only maternal origin.

The total number of COs per autosomal arm segregating in the pedigree ranged between 0 and 11 (mean: 6.59) when considering both paternal and maternal COs, between 0 and 7 (mean: 3.32) considering only paternal origins, and between 0 and 6 (mean: 3.26) considering only maternal origins (**Figure 2B-D; Table S4**). This corresponds to sex-average recombination distances for the autosomal arms ranging between 0 and 91.7 cM (mean: 54.9 cM), paternal recombination distances between 0 and 116.7 cM (mean: 55.4 cM), and maternal recombination distances between 0 and 100 cM (mean: 54.4 cM). The recombination distances between paternal and maternal chromosome arms did not differ significantly (paired t-test; t = -0.21, df = 33, p = 0.84). For the Z chromosome, the recombination distance was 8.3 cM for the PAR (sex-average recombination) and 100 cM for the remaining part (paternal recombination).

The sex-average recombination rate for the autosomal arms ranged between 0 and 17.7 cM/Mb (median: 3.06 cM/Mb), whereas the paternal rate varied between 0 and 21.2 cM/Mb (median: 2.93 cM/Mb), and the maternal rate between 0 and 17.7 cM/Mb (median: 2.73 cM/Mb) (**Figure 3A**). There was no significant difference in recombination rate between the sexes (paired t-test; t = -0.16, df = 33, p = 0.88). There was a pronounced non-linear negative association between the recombination rate and the size of the chromosome arms (**Figure 3B**). For the Z chromosome, the recombination rate was 7.58 cM/Mb for the PAR and 1.16 cM/Mb for the remaining part of the chromosome.

**Figure 3.**
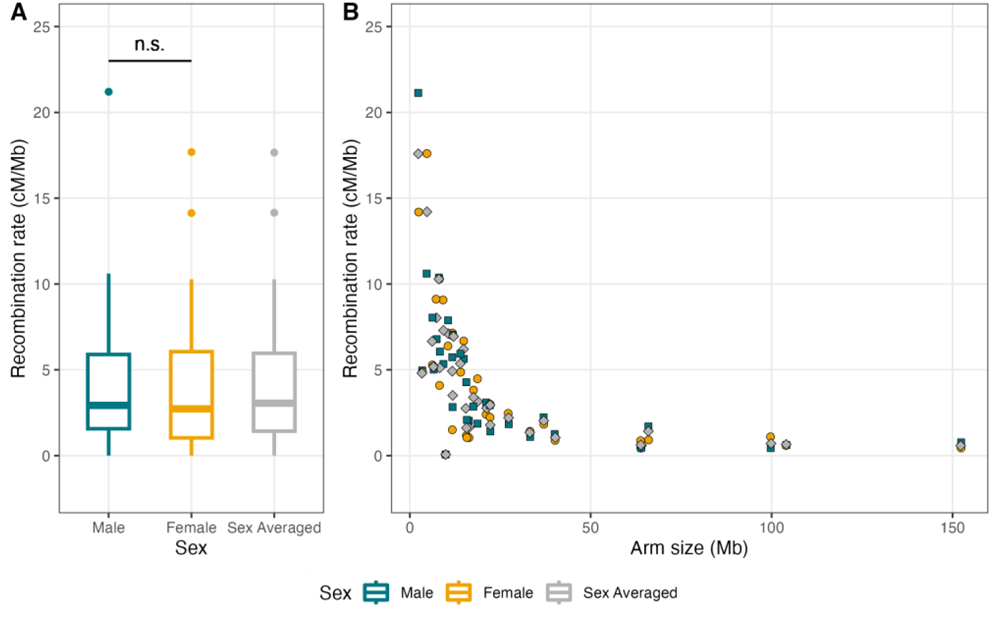
(A) The distribution of male, female and sex-averaged recombination rates on autosomal arms. (B) The association between the male (green rectangles), female (yellow circles) and sex-averaged (grey diamonds) recombination rates and the size of autosomal arms.

The position of the 224 autosomal COs showed strong bias towards the telomeric end of the autosomal arms, and this was true for both paternal and maternal chromosome arms (**Figure 4A**). However, this bias was significantly more pronounced on paternal chromosomes, both when considering the physical distance from the telomeric end (male median position: 1.57 Mb; female median position: 2.10 Mb; Mann-Whitney U test; U = 7732, p = 0.003; **Figure 4B**) and the proportional distance on the chromosome arms (male median position: 9.29%, female median position: 13.14%, Mann-Whitney U test; U = 7365, p = 0.024; **Figure 4C**).

**Figure 4.**
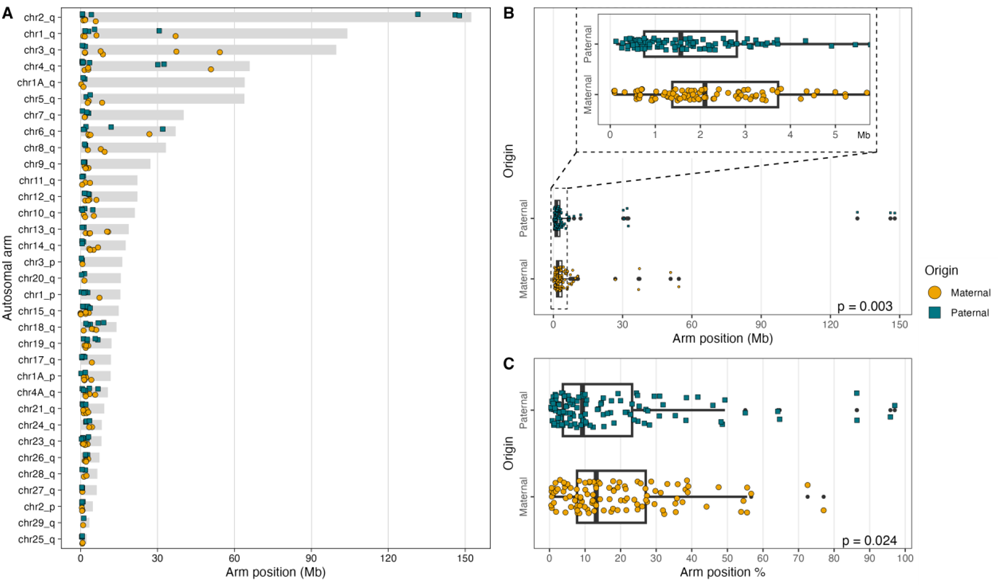
The bias of COs towards the telomeric ends of chromosomes. (A) The location of CO positions of paternal and maternal origins on each chromosome arm. (B) The physical distance of CO positions from the telomeric end of autosomal arms. (C) The proportional distance of CO positions from the telomeric end of autosomal arms. Grey bars (A) indicate the sizes of the autosomal arms. The colouration and shape of points (A-C) indicate paternal (green squares) and maternal (yellow circles) CO events.

Regarding the distribution of autosomal COs in relation to gene features, 109 COs occurred in intergenic regions and 115 in genic regions, which is not significantly different from an expectation based on the size of these regions (chi-square test; χ^2^ = 2.77, df = 1, p = 0.096; **Figure 5A**). Out of the 115 autosomal COs in genic regions, 91 were in introns and 24 in exons, which is significantly different from an expectation based on the size of these regions (chi-square test; χ^2^ = 3.96, df = 1, p = 0.047). Note that the data suggest a relatively higher frequency of COs in exons than in introns as exons are much less abundant in the genome than introns. Among the 24 COs in exons, 17 occurred in CDS and 8 in UTRs, which is not significantly different from an expectation based on the size of these regions (chi-square test; χ^2^ = 1.34, df = 1, p = 0.25). Regarding the distance to the closest genes, 90% of the autosomal COs were located between -34 kb (upstream) and 47 kb (downstream) to the closest genes (**Figure 5B**).

**Figure 5.**
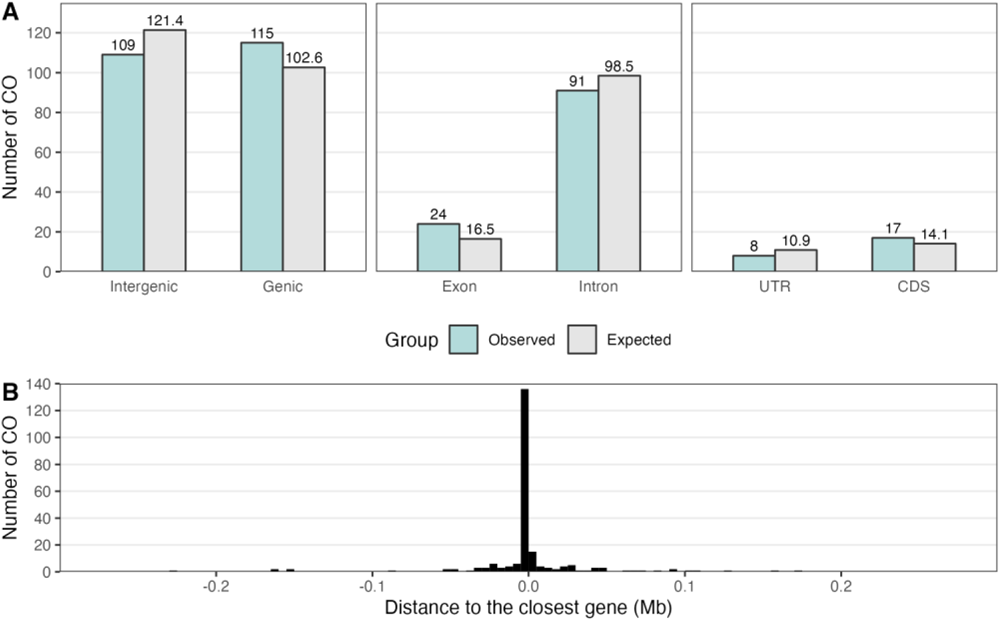
The relationship between autosomal CO events and gene features. (A) Number of COs in intergenic and genic regions, in exons and introns, and in UTR and CDS (green bars). Aslo given are the expected numbers based on the size of these gene features within 6 Mb of the end of the chromosome arms (grey bars). (B) The distribution of the distance to the closest gene for autosomal CO events.

## 4 Discussion

Quantifying the recombination landscape and identifying chromosome regions with varying rate of recombination can improve our understanding of the evolutionary impact of gene linkage. Additionally, it can provide insights into the mechanisms underlying the pairing and segregation of homologous chromosomes during meiosis. In this study, we analysed whole-genome sequencing data from a three-generation pedigree of the great reed warbler to precisely locate crossover (CO) events. The high resolution of our analysis enabled us to examine the relationship between COs and various chromosomal features, as well as compare the recombination patterns between paternal and maternal chromosomes.

We observed seven CO events on the Z chromosome, one in the pseudoautosomal region (PAR) of maternal origin, and six in the remaining part where only males recombine. The recombination pattern on the Z chromosome was generally similar to that of similarly sized autosomes, *i.e.*, concentrated towards the chromosome ends. However, since the sex chromosomes segregate differently in males and females, and since recombination only occurs in the PAR in females, we will not further discuss the few cases of COs on the Z chromosome.

Our analysis focused on 408 autosomal arms segregating in the pedigree (34 arms; 12 meioses per arm among the 6 offspring), and we identified 224 COs. The majority of autosomal arms had either zero or one CO, with only nine cases of two COs and one case of three COs on a single chromosome arm being detected. This implies that 52.2% of the autosomal arms had at least one CO. This finding aligns with two expectations: (i) the probability of a CO product being passed down to the offspring is 50% per chiasma, considering the presence of two sister chromatids with one having a CO, and (ii) the hypothesis of an obligate chiasma requirement, suggesting that at least one chiasma per chromosome arm is necessary for proper segregation of homologous chromosomes (Coop and Przeworski 2007; Stapley et al. 2017). It is worth noting that the terms “crossover” and “chiasma” are sometimes used interchangeably, but that we refer to CO as the genetic recombination event and to chiasma as the cross-shaped interaction between non-sister chromatids of homologous chromosomes during meiotic prophase. The obligate chiasma requirement indicates the necessity to connect homologous chromosome arms, enabling their alignment on the spindle in metaphase I, and thereby facilitating correct segregation (Petronczki et al. 2003). However, there are exceptions to this requirement, most obviously among species exhibiting achiasmy, where chiasmata are absent in one of the sexes (Haldane 1922; Huxley 1928; John et al. 2016). Furthermore, investigations into the obligate chiasma requirement in mammals using cytogenetic and phylogenetic methods identified multiple independent shifts from one chiasma per chromosome arm to one chiasma per chromosome across the phylogenetic tree, extending the hypothesis to a minimum of one CO per chromosome (Dumont 2017). In this study of the great reed warbler pedigree, we detected at least one CO at all chromosome arms, except for chromosome arm 4p (the short arm; **Table S4**) where we did not detect any CO. This could be due to chance, as zero COs can be expected to occur among 12 meiotic events (one parental and one maternal meiosis for each of the six offspring). Alternatively, it may indicate incomplete assembly of the telomeric end of this chromosome arm, leading to missed CO events, or that the location of the centromere is more telocentric than we estimated from the low nucleotide diversity and its location in the zebra finch. The limited occurrence of chromosome arms with multiple COs in the great reed warbler (ten cases of 213 autosomal arms with at least one CO) supports the notion that CO interference reduces the likelihood of additional COs (Sturtevant 1915; Muller 1916; Otto and Payseur 2019). Notably, recombination interference can be positive, resulting in fewer and/or more spaced COs (Coop and Przeworski 2007) as seems to be the case in great reed warblers, or negative, leading to a higher recombination rate than expected, which is observed in some plants and animals (Auger and Sheridan 2001; Aggarwal et al. 2015).

A study of recombination in humans did not only confirm the obligate chiasma requirement but also emphasised the significance of proper location of chiasmata for accurate segregation of homologous chromosomes during meiosis (Coop and Przeworski 2007). Our analysis revealed an extreme bias of COs towards the sub-telomeric regions of the chromosomes, with approximately 87% of COs occurring within approximately 6 Mb from the chromosome ends. This particular pattern has been observed in various other taxa (Sardell and Kirkpatrick 2020), including in zebra finch and collared flycatcher (Backström et al. 2010; Smeds et al. 2016), indicating a widespread phenomenon. The underlying reason for the bias of COs towards sub-telomeric regions remains an open question. One possibly is that the heterogeneous distribution of COs on chromosomes is regulated by specific DNA sequence structures or motifs. In humans, an overrepresentation of a 13-mer DNA motif has been identified as a factor inducing recombination, with the motif being recognised and bound by the zinc finger of PRDM9, a histone methyltransferase (Myers et al. 2008). While the *PRDM9* gene is present in many mammals, it is absent in dogs and birds, suggesting that alternative mechanisms exist (Baudat et al. 2010; Paigen and Petkov 2018). Other genes such as *RNF212*, *CPLX1* and *REC8* have been repeatedly implicated in recombination events in mice, humans, cattle and sheep (Reynolds et al. 2013; Kong et al. 2014; Ma et al. 2015; Johnston et al. 2016). A search for DNA motifs associated with recombination in estrildid finches identified a few candidate motifs, but none were found to be causative (Singhal et al. 2015). Likewise, we did not find evidence of overrepresented motifs around the recombination positions that we detected in the great reed warbler (H. Zhang *et al*., unpublished). Regarding the location of COs in relation to genes, we found that the frequency of COs was similar between intergenic and genic regions. However, within genes, COs occurred more commonly in exons than in introns, suggesting a potential influence of gene structure of recombination events. These patterns differ from those in human and *Drosophila* where recombination rates are generally lower within exons compared to introns and intergenic regions (Kong et al. 2010; Miller et al. 2012).

The number of recombination events was similar between the sexes at both the chromosome and genome level in the great reed warbler, and both sexes exhibited a strong bias towards COs located near the telomeric ends of chromosomes. However, we observed that COs occurred significantly closer to the telomeres on paternal chromosome arms compared to maternal chromosome arms. Previous studies on the great reed warbler, which relied on a limited set of markers but a large multi-generation pedigree, reported approximately twice as high recombination rate in females compared to males (Hansson et al. 2005; Dawson et al. 2007). The discrepancy between these findings and our study likely arises from the fact that the markers used in the former studies were primarily located in the central regions of chromosomes, resulting in an underrepresentation of sub-telomeric CO events, which are biased towards males as observed in the present study. This emphasises the importance of thoroughly screening the entire length of chromosomes to accurately characterise potential sex biases in recombination. In a separate project, we have conducted genotyping using genome-wide distributed restriction site-associated DNA (RAD) markers on a multi-generational pedigree (Ponnikas et al. 2022). Although this dataset had a smaller number of markers (approximately 50k SNPs) compared to the present study (approximately 5M SNPs), it included multiple males and females. Preliminary analyses of the autosomal RAD data set confirmed that recombination is biased towards the end of chromosomes in both sexes, and that females have a higher frequency of recombination in the central regions of chromosomes compared to males (S. Ponnikas *et al*., unpublished). Furthermore, male-biased recombination in sub-telomeric regions has been observed in several other taxa including in birds (Backström et al. 2010; Smeds et al. 2016; Sardell and Kirkpatrick 2020), while female-biased recombination around centromeres or centrally on chromosomes has been reported in some cases (Venn et al. 2014; Sutherland et al. 2017; Sardell and Kirkpatrick 2020). Understanding the mechanisms underlying sex differences in the recombination landscape will require considering the combined effects of sex-specific centromeric and telomeric influences, as well as how telomere-guided initiation of recombination clusters COs in sub-telomeric regions in both sexes (Higgins et al. 2012; Haenel et al. 2018; Otto and Payseur 2019).

In conclusion, recombination plays a crucial role in evolution by generating new haplotypes that natural selection can act upon. In this study, we utilised whole-genome sequencing data of a three-generation pedigree of the great reed warbler to locate CO positions with high precision and investigate sex-specific patterns of recombination. We found that the overall number of recombination events was similar between the sexes. However, when examining the distribution of CO positions, we discovered a pronounced bias towards the telomeric ends of the chromosomes in both sexes, with a particular strong bias on parental chromosomes. We also found that intra-genic COs were more frequently located within exons than in introns. Elucidating the sex-specific CO landscape in the great reed warbler provides valuable evidence to gain deeper understanding of recombination, a key mechanism in shaping the genetic diversity within populations.

## Data Availability Statement

All sequence data used for this study are accessible under BioProject ID’s PRJNA970100.

## Acknowledgements

Sample collection was granted with the appropriate permissions from the Linköpings Djurförsöksetiska nämnd (Dnr 36-11, Dnr 44-14 and ID1633). Sequencing was performed by the SNP&SEQ Technology Platform at Uppsala Genome Center, which is part of National Genomics Infrastructure (NGI) Sweden, and Science for Life Laboratory (SciLifeLab) supported by the Swedish Research Council (and its Council for Research infrastructure, RFI) and the Knut and Alice Wallenberg Foundation. Bioinformatics analyses were performed on computational infrastructure provided by the Swedish National Infrastructure for Computing (SNIC) at Uppsala Multidisciplinary Center for Advanced Computational Science (UPPMAX). Field work was supported by the Kvismare Bird Observatory (report no. xxx).

## Author contributions

H.Z., D.H. and B.H. designed and planned the study. H.Z. performed the laboratory work of preparing genomic DNA. H.Z. performed the bioinformatic analyses with input from M.L and B.H. H.Z. and B.H. wrote the manuscript with input from M.L., S.P., and D.H. All authors approved the final version of the manuscript.

